## Supplementary information for "SEMORE: SEgmentation and MORphological fingErprinting by machine learning automates super-resolution data analysis"

### Supplementary Tables

**Supplementary Table 1**: Description of each morphology fingerprint feature.

| # | FEATURE | SUB-SET | DESCRIPTION |
|---|---|---|---|
| 1 | x ratio | Symmetry | Ratio of points between the left and right side of the y-axis |
| 2 | y ratio | Symmetry | Ratio of points between above and below the x-axis |
| 3 | ratio sym | Symmetry | x ratio * y ratio |
| 4 | x dist ratio | Symmetry | Ratio of longest absolute distance compared to total span (x-axis) |
| 5 | y dist ratio | Symmetry | Ratio of longest absolute distance compared to total span (y-axis) |
| 6 | Pearson | Symmetry | Pearson coefficient. |
| 7 | Spearman | Symmetry | Spearman coefficient |
| 8 | Area | Geometric | Area estimation found through accumulation on triangle areas. |
| 9 | Density | Geometric | Points in aggregate divided with area. |
| 10 | Cut distance | Geometric | Longest allowed distance for pairwise connection |
| 11 | Mean k | Graph network | Average amount of neighbors/connection |
| 12 | Median k | Graph network | Median of neighbors/connection |
| 13 | K max | Graph network | Max of neighbors/connection |
| 14 | L_s_d | Graph network | Distance between most separated points |
| 15 | L_s_path | Graph network | Distance along minimum spanning tree between most separated points |
| 16 | L_s_step | Graph network | Number of bridges used in L_s_path |
| 17 | L_s_mean | Graph network | Average distance connection in L_s_path |
| 18 | L_s_median | Graph network | Median of distance connection in L_s_path |
| 19 | L_l_max | Graph network | Max distance connection in L_s_path |
| 20 | L_s_effectiveness | Graph network | L_s_d divided with L_s_step |
| 21 | L_s_ratio | Graph network | Ratio between the L_s_d and L_s_path |
| 22 | L_l_path | Graph network | Longest possible one-way road through the minimum spanning tree |
| 23 | L_l_d | Graph network | Direct distance between start and end points in L l path |
| 24 | L_l_step | Graph network | Same as 14 now with L_l_path |
| 25 | L_l_mean | Graph network | Same as 15 now with L_l_path |
| 26 | L_l_median | Graph network | Same as 16 now with L_l_path |
| 27 | L_l_max | Graph network | Same as 17 now with L_l_path |
| 28 | L_l_effectiveness | Graph network | Same as 18 now with L_l_path |
| 29 | L_l_ratio | Graph network | Same as 19 now with L_l_path |
| 30 | Ls_ll_d ratio | Graph network | Ratio between L_s_d and L_l_d |
| 31 | Ls_ll_path ratio | Graph network | Ratio between L_s_path and L_l_path |
| 31+N | Mu 1-N | Graph network | Mean of N fitted Gaussians over connections (k) |
| 31+2N | Sig 1-N | Graph network | Standard deviation of N fitted Gaussians over connection (k) |
| 31+3N | W 1-N | Graph network | Weights of N fitted Gaussians over connection (k) |
| 32+3N | State diff | Graph network | Average difference between Gaussians |
| 33+3N | Variance | Circularity | Variance of distance from the core to morphology edge |
| 34+3N | Circularity | Circularity | $4 \cdot \pi \cdot area \,/\, circumfrence^2$ |
| 35+3N | Convexity | Circularity | The area of convex hull dived with area of contour line |
| 36+3N | Circ inertia | Circularity | Assuming ellipse, ratio between vertex |
| 37+3N | Sph value | Circularity | combination of the 4 circularity features E[(1 −V ar) + Circ + Conv + Inert] |

### Supplementary Figures

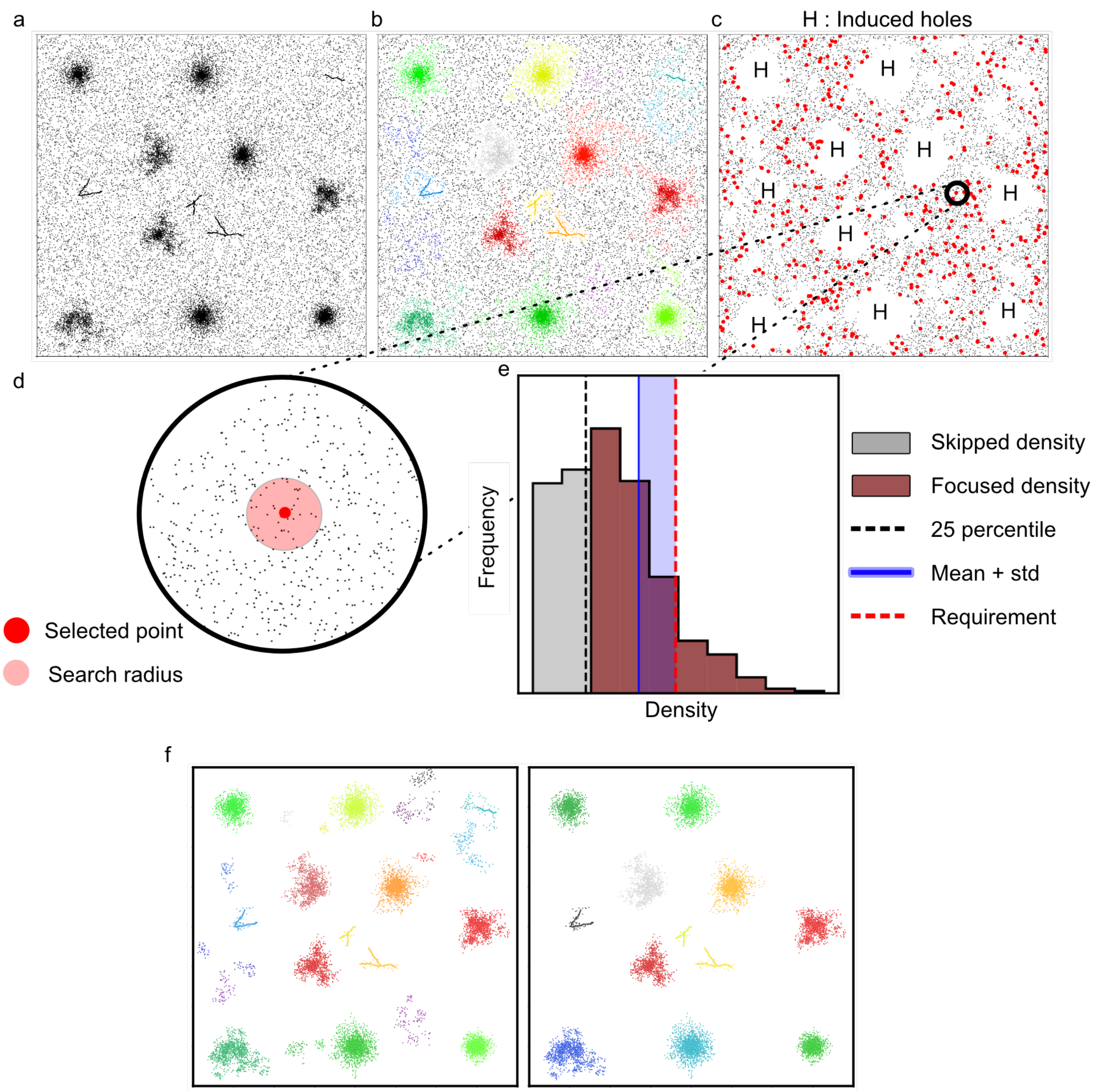


**Supplementary Fig. 1: Complete pipeline and resulting effect of introduced smart density filter. a,** Raw simulated data with an experimental relevant noise ratio. **b,** the initial clustering with noise in black and the high-density areas in colors. **c,** The initial found noise, with the high-density areas removed, resulting in multiple holes (denoted "H"). 500 noise points (red) are randomly selected for noise density estimation. **d,** A zoom-in of a selected noise point. For each selected point a search radius of 0.03 (in standardized space) is drawn from which the density is calculated. **e,** The accumulated densities in a histogram. Due to the holes present in **c,** some selected points may experience sparse densities, to account for this, a 25-percentile mask is applied (grey) to avoid any underestimation. This results in focused densities (red-brown) from which a minimum density requirement is then calculated as the mean + the standard deviation. **f,** Comparison visualization, the presented data set treated through SEMORE without the smart density filtering, clearly extracting multiple non-aggregate structures, while when applied completely removing these structures, drastically lowering the overall false positive rate.

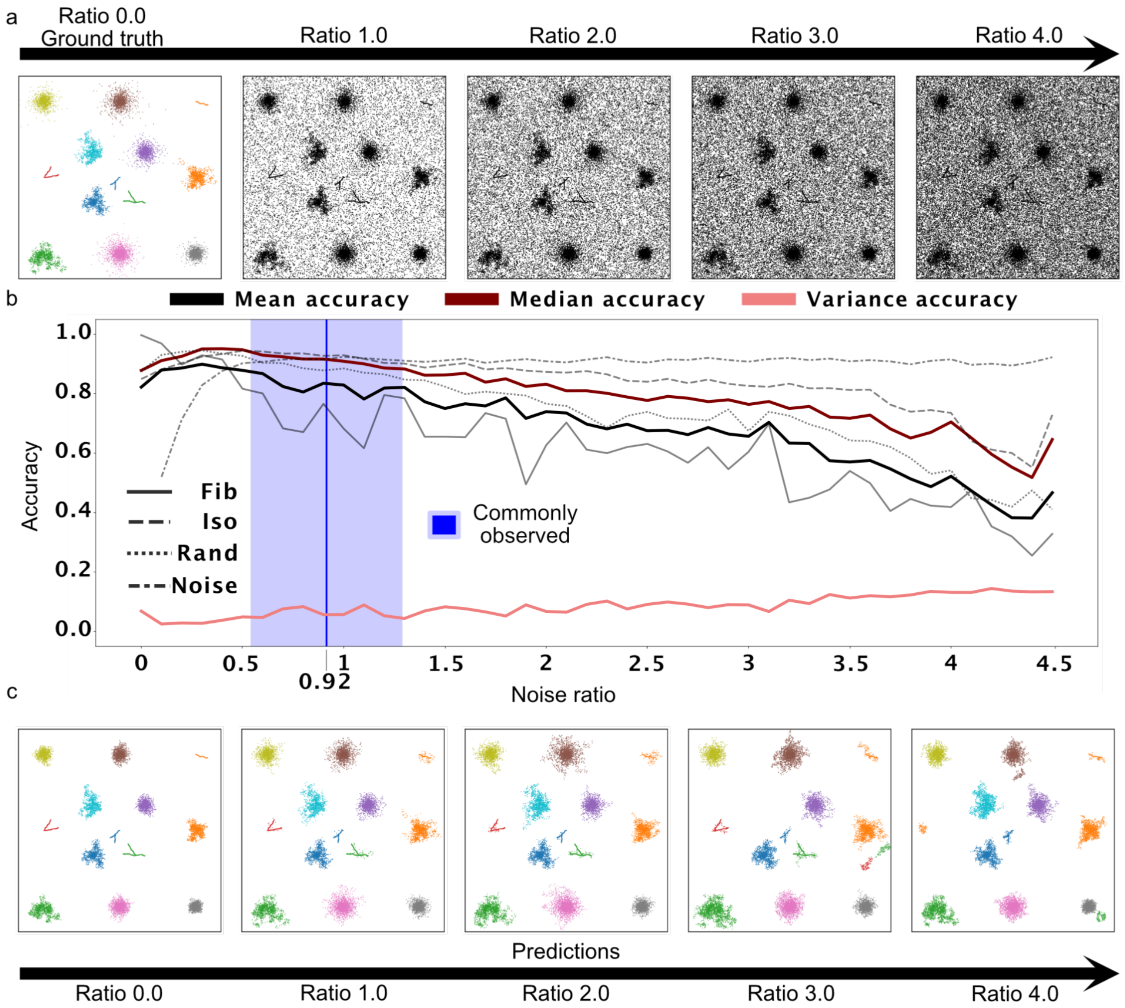


**Supplementary Fig. 2: SEMORE out-of-box noise stress test of non-overlapping highly diverse aggregates. a,** 5 experiments were simulated without noise, each containing 13 aggregates of randomly selected simulation types. To ensure minimal overlapping, each aggregate was kept in the same grid throughout the 5 experiments. The noise was incremented for each experiment with a 0.1 ratio (N_noise / N_label) up to a 4.5 noise ratio. **b,** SEMORE accuracy (see Methods) against noise ratio increase. Both mean (black) and median (dark red) start with a positive slope and reach their highest performance at 90% and 95% respectively for a noise ratio of ~0.4, which corresponds to the general assumption of an unavoidable noise present in the training sets (grey dash-dot). Further noise ratio increases results in a linear decrease in the accuracy. Fibril (gray solid) displays the steepest decline, due to its smaller spatial occupation and density compared to isotropic (gray dashed) and random (gray short dashed). At an unreasonably high noise ratio of noise 4.5, the mean and median were 38% and 52% respectively with a variance of around 15%. The biologically relevant and experimentally recorded noise of 0.92 is displayed with a blue line with a spread of 0.37 shown as blue transparency on both sides. **c,** Predicted classification labels for the noise ratios. SEMORE generally separates noise from labels for experimentally relevant noise (<1). At higher noise density, the predictions include more false positive locations and whole aggregates disappearing. In the experimentally relevant noise range, SEMORE correctly outputs the underlying structures at median accuracy of up to 93%.

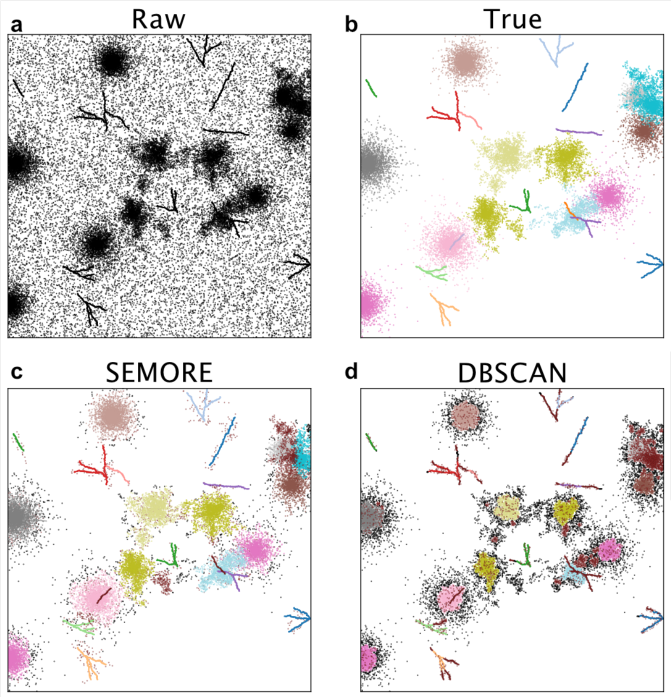


**e**

| Single aggregate wise | SEMORE | DBSCAN |
|---|---|---|
| Accuracy median | 89.5 % | 70.1 % |
| Accuracy mean | 72.0 % | 57.1 % |
| Accuracy Var | 12.0 % | 11.7 % |
| Noise separation | 89.3 ±2.3 % | 74.5 ± 3.1 % |

**f**

SEMORE

| Type | Precision | Recall | F1 |
|---|---|---|---|
| Weighted | 86.5 ± 5.1% | 88.6 ± 3.2% | 87.2 ± 4.4% |
| macro | 74.0 ± 7.1% | 80.6 ± 7.2% | 76.4 ± 7.2% |
| micro | 87.9 ± 4.8% | 87.5 ± 5.0% | 87.7 ± 4.8% |

DBSCAN

| Type | Precision | Recall | F1 |
|---|---|---|---|
| Weighted | 76.4 ± 7.2% | 75.4 ± 4.1% | 73.5 ± 5.7% |
| macro | 65.3 ± 9.5% | 64.3 ± 7.3% | 63.5 ± 8.3% |
| micro | 80.0 ± 8.0% | 67.9 ± 6.0% | 73.5 ± 6.7% |

**Supplementary Fig. 3: Comparison of operational performance of segmentation module in stress test containing simulated overlapping protein aggregates where temporal information is included. a,** Representative visualization of the raw data based on simulations containing all three simulation-aggregation types resulting in a highly dense and overlapping aggregate diverse system. **b,** True label illustration without noise. **c, d,** Segmentation by our method and DBSCAN. Wrongly labelled aggregates are colored brown and wrongly predicted noise is colored black. Both methods' hyperparameters were optimized once for the same simulated experiment and were used for multiple simulations. DBSCAN misses or incorrectly classifies aggregates in proximity and when structures overlap. **e,** Accuracy measure (for 50 experiments containing 25 aggregates each totaling 1250 aggregates) defined as TP/ (TP +FP +FN). As this measure is done for each individual aggregate, the accuracy ranges between 0 and 1 for all structures removing the bias imposed by the large aggregate size variations. SEMORE achieved 89.5% median accuracy, 19.4% higher than DBSCAN with only a 0.3% increase in variance. Furthermore, SEMORE achieves a higher noise separation as well, resulting in both more and cleaner aggregates. **f,** Performance tables for both models with all metrics extracted per experiment. Due to the high diversity of the experiments multiple evaluation approaches are reported: Weighted being evaluation metric per label weighted by TP per label. Macro being the metric for each label unweighted. Micro being total TP, FN and FP combined to a global metric. In general, SEMORE outperforms DBSCAN mainly due to its lack of ability to utilize the temporal element, although the temporal dimension was provided to DBSCAN, as well as the addition of density smart filtering.

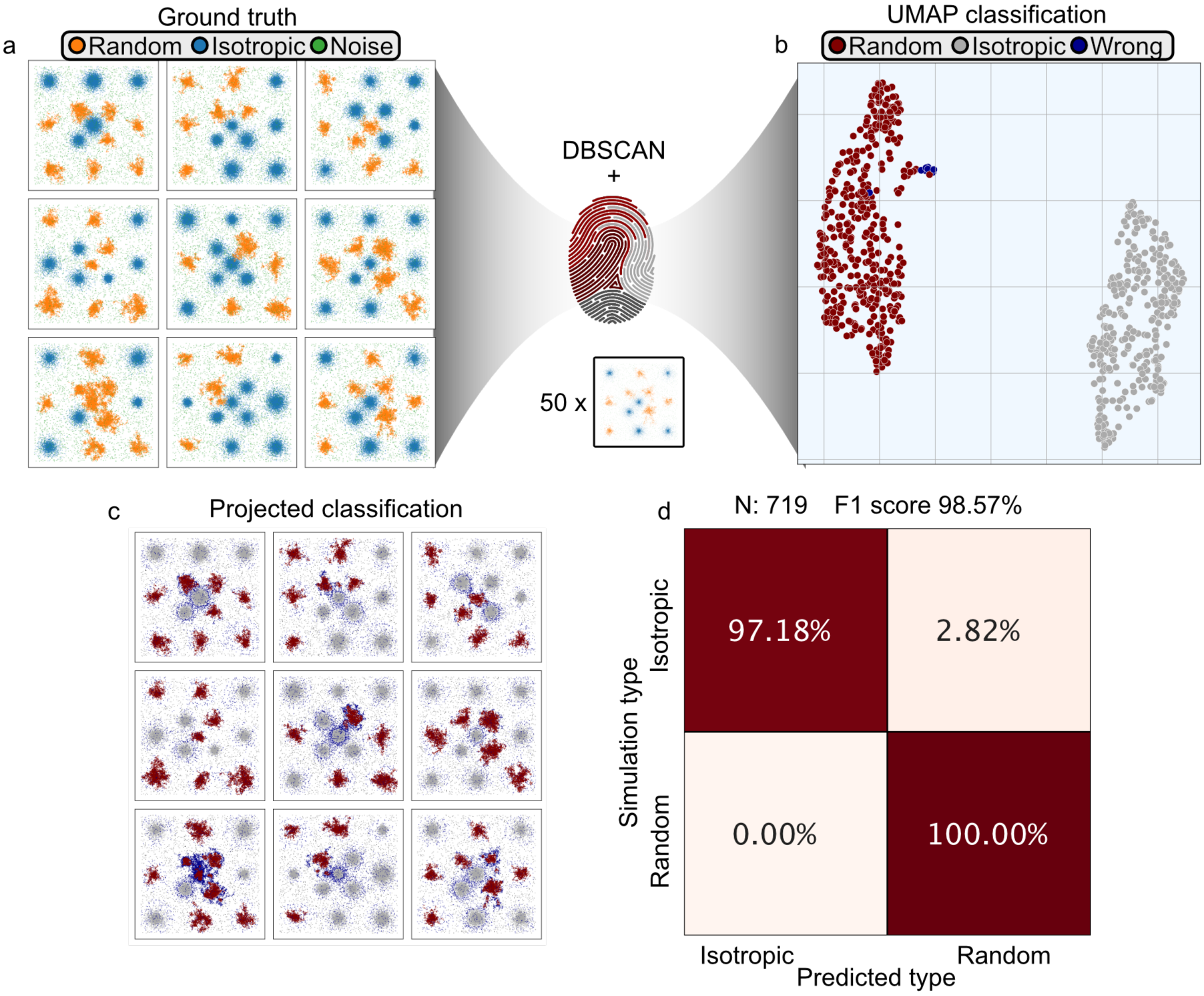


**Supplementary Fig. 4: Classification capabilities of unsupervised morphology fingerprinting, in high-density regions of diverse aggregate morphologies for SMLM without temporal features. a,** 9 representative simulated experiments of the total 50, each containing 13 aggregates with the same start seed location throughout the simulations. The aggregate type is randomly selected as either isotropic or random and colored accordingly. **b,** To mimic an analysis pipeline of SMLM data without temporal resolution, each simulation is treated through a DBSCAN (eps = 500, min_sample = 25) to segment the contained aggregates. The SEMORE fingerprint module is then used to extract the morphology fingerprint for each of the extracted structures. The resulting features are then embedded through a UMAP ( n_neighbors = 15, min_dist = 0.1) and clustered in the embedded space through another DBSCAN (eps = 1, min_sample = 10). This results in 2 clearly separated clusters used to predict the dominant aggregate type contained within, with Random colored dark red, Isotropic colored gray and wrong prediction colored blue. **c,** The corresponding prediction colored on the raw data, for visual comparison while still coloring incorrect prediction as blue. **d,** Confusion matrix for the binary prediction of aggregation type, achieving an F1 score of 98.57% indicating almost perfect predictions and information capture within the fingerprint.

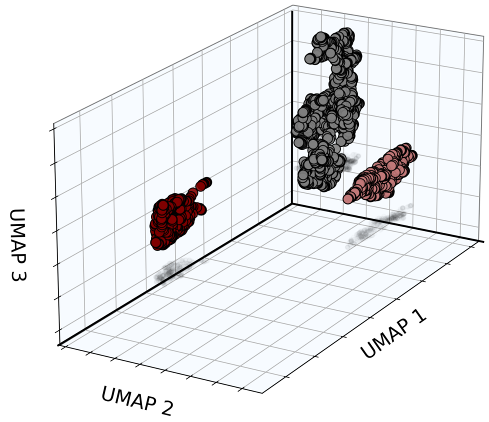


**Supplementary Fig. 5**: **Resulting UMAP from SEMORE clustering applied on the simulated structures with the smart density filtering.** This results in 3 separate clusters colored according to the color mapping as seen in Fig. 2b. However, as the smart density filtering removes the “noise”-aggregate the noise cluster is not present in the UMAP, which is a testimony to the utility of the smart density filter.

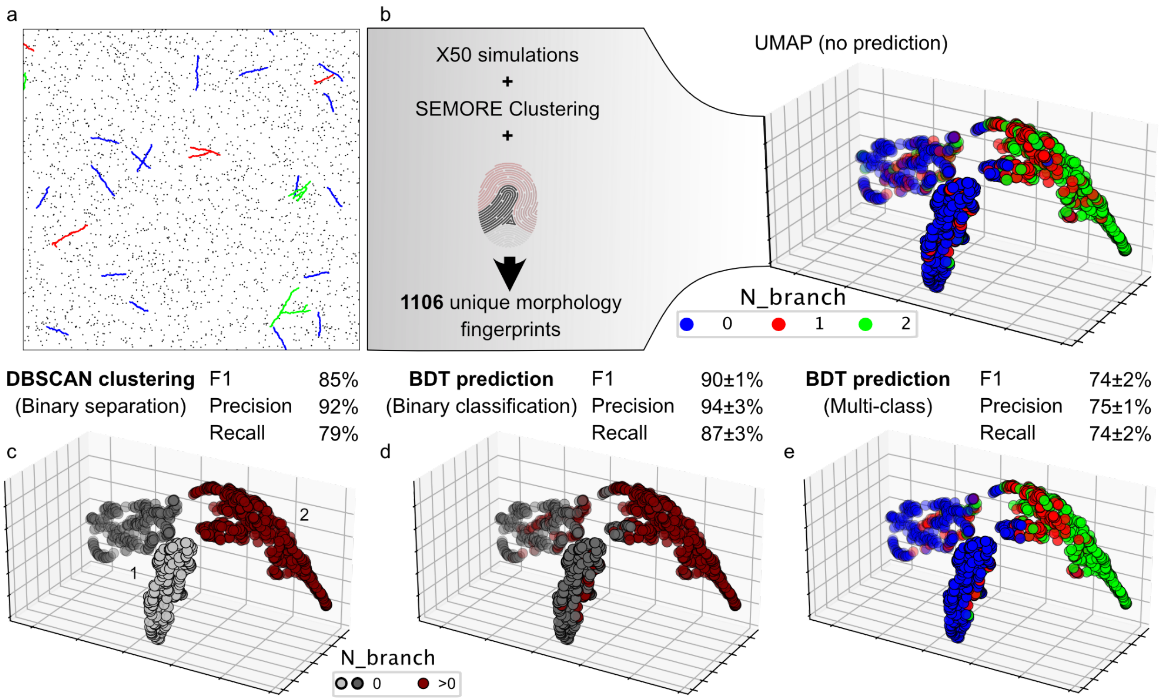


**Supplementary Fig. 6: Fibril branch classification performance through feature-class specific investigation. a,** Representative examples of the simulated experiments with the fibrils colored according to their number of branches. **b,** Treating the 50 simulated experiments results in 1106 unique fibril morphology fingerprints. These were embedded through UMAP (n_neighbors = 15, min_dist = 0.1, n_components = 3) and each point colored to its corresponding aggregate branching degree. **c,** A DBSCAN (eps = 1, min_sample = 10) clustering performed in the embedded space for binary classification of non-branching (positive) and branching (negative) fibrils. The results yielded three distinct clusters: Gray clusters being highly dominated by non-branching fibrils and collectively representing cluster 1, whereas cluster 2 in red is highly dominated by branching. This binary classification results in an overall F1 score of 85%, with a recall of 79% and precision on 92 %. **d,** To demonstrate the versatility of the morphology fingerprinting, a boosted decision tree (BDT) was fitted and evaluated through a k = 5-fold, achieving a mean F1 score of 90±1% which increased to 94±2% when exposing all features of the morphology fingerprint. **e,** Furthermore, the same BDT model was fitted to classify the multi-class problem of N_branches, resulting in a mean F1 score of 74±2% increasing to 82±3% with the use of all features. The overlap of classes seen in the embedded space can be caused by multiple factors, as the investigated feature class only contains information regarding morphology. This results in the blinding of fibril outliers, e.g. late branching as seen in red aggregates in **a,** or highly diverse angle distribution in back-bone growth. This would also explain the low but prominent increase in BDT performance through all the features compared to only circularity which comprehensively contains 5 out of 40+ features.

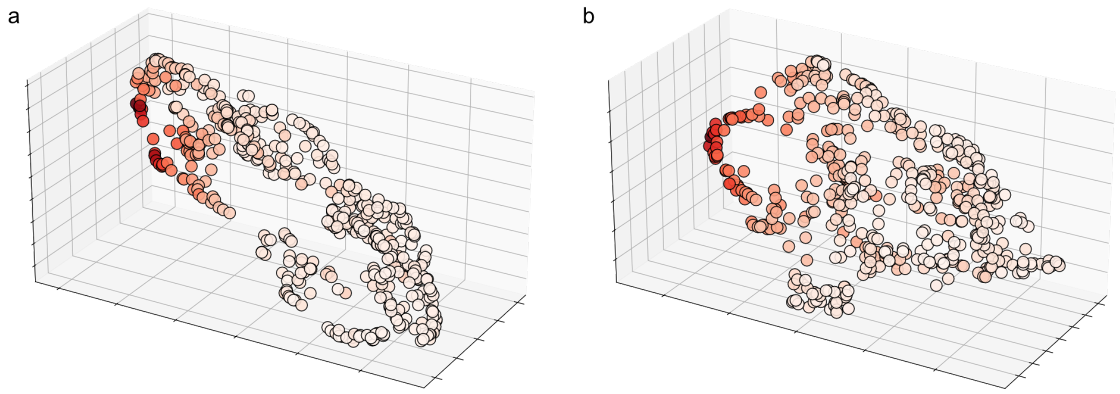


**Supplementary Fig. 7: Simulation type specific UMAP investigation based on circularity feature class. a,** Corresponding embedded features from simulated isotropic aggregates, the general structure of the embedding depicts a linear dependency on the aggregate morphology due to no fundamental changes in growth pathways, as compared to the branching behavior in fibrils. The comparably narrow spread in points is due to the segmentations not being perfectly extractions and the high diversity in size. **b,** The embedding for simulations based on steric hindrance ("random" aggregate) has a wider spread out due to stochasticity in their s growth kinetics and morphology. Both the UMAP and features used are the same as seen in Supplementary Fig. 5, with the datapoints color corresponding to the variance-feature value, with darker color being a higher value.

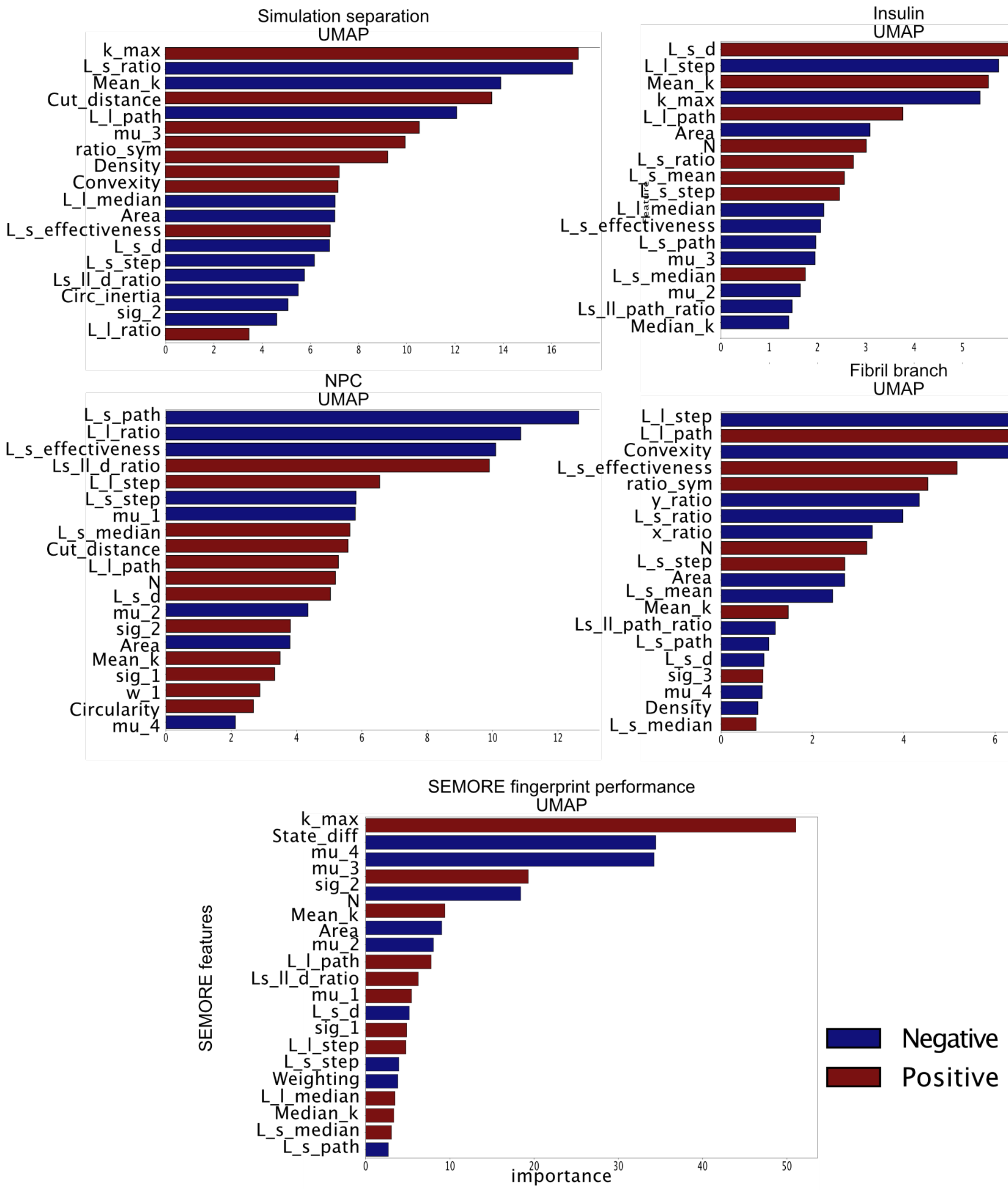


**Supplementary Fig. 8:** Extracted feature importance of all UMAPs presented in the article, the bars are colored according to the weighting (positive: red, negative: blue) and the importance values are found through linear discriminant analysis (LDA) fitted to the predicted labels.

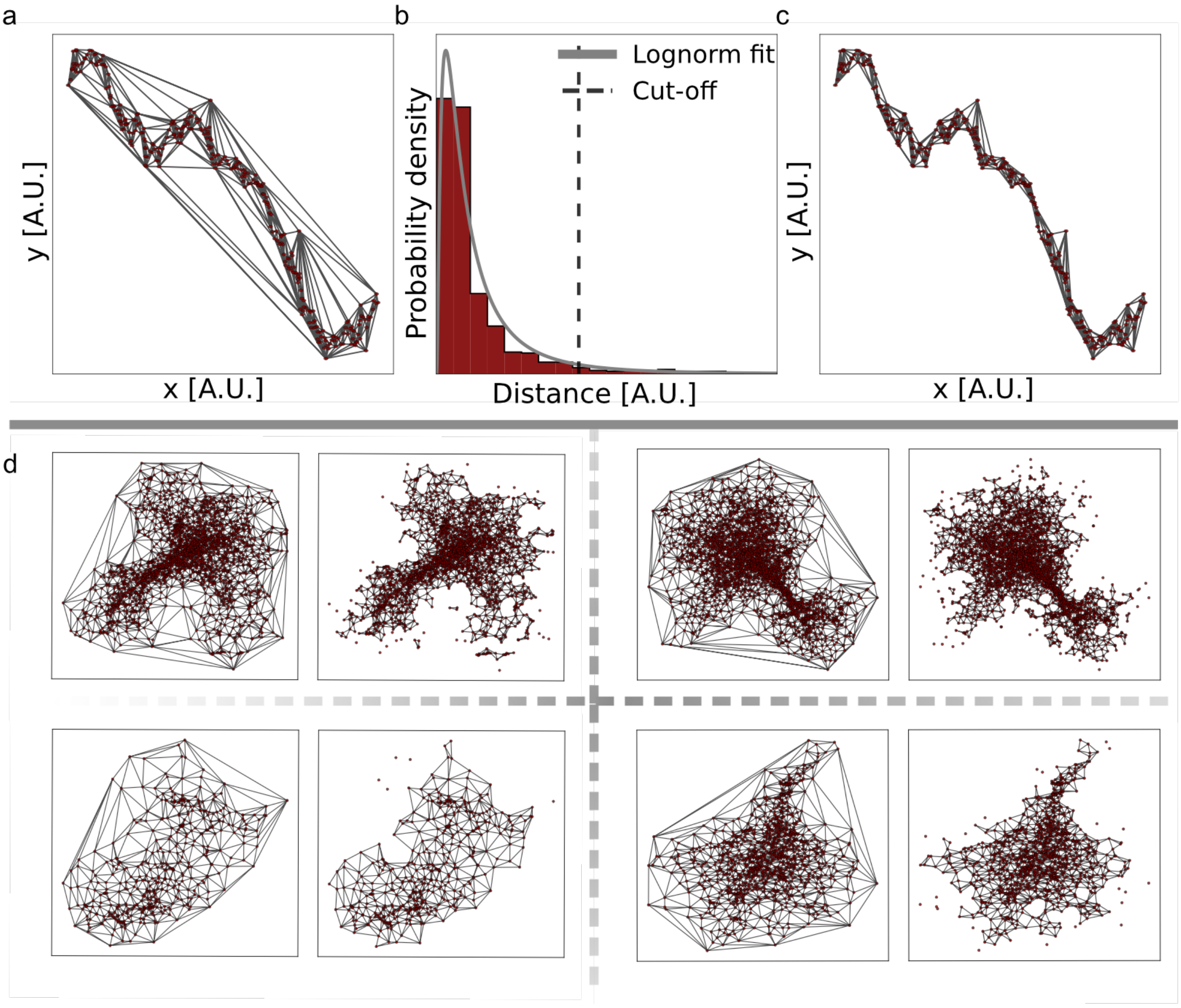


**Supplementary Fig. 9: Demonstration and visualization of structure-polygon for size estimation. a,** Displaying a simulated fibril structure, along the collected localizations (dark red) and Delaunay triangulation (grey). **b,** The triangle-edge distance distribution is fitted by the lognormal. From this, a distance threshold is calculated as the right-tail 5% probability or below. **c,** The threshold is then used to prune the triangle edges, from which only the remaining closed triangles are used to calculate the area of the given structure. **d,** Shows the Delaunay triangulation along its pruned counterpart from which the area can be evaluated, both as a static image or temporally as the structure grows.

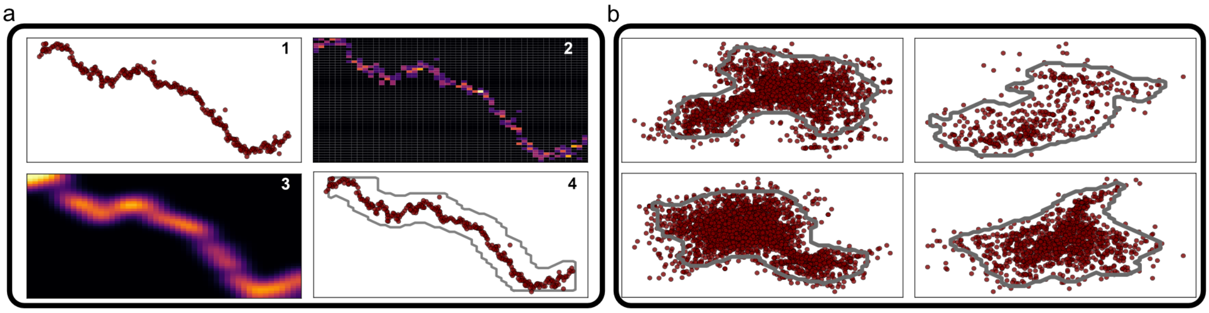


**Supplementary Fig. 10: Pipeline for morphology edge estimation. a,** The 4-step process for achieving the circumference estimate of a given structure a.1) The x, y coordinates of the investigated structure are collected, 2) the data points are fed into a 2-dimensional 50x50 bin histogram, 3) a Gaussian blur kernel convolved over the counts in the histogram to create a smoother structure. 4) From a pre-defined? percentile threshold (70% default), a binary mask is acquired and extracted through a one-level contour analysis. This contour can then be used along with the pre-defined core to calculate the corresponding features of the circularity sub-set. For fine-tuning, the number of bins, sigma of the Gaussian kernel and the percentile cut-off are adaptable, while for a density-dependent circumference, the amount of connection acquired from the graph network can be fed into the initial histogram as weights (a.2). **b,** For clarification the method is visualized for the same 4 random insulin aggregates as in Supplementary Fig. 9.

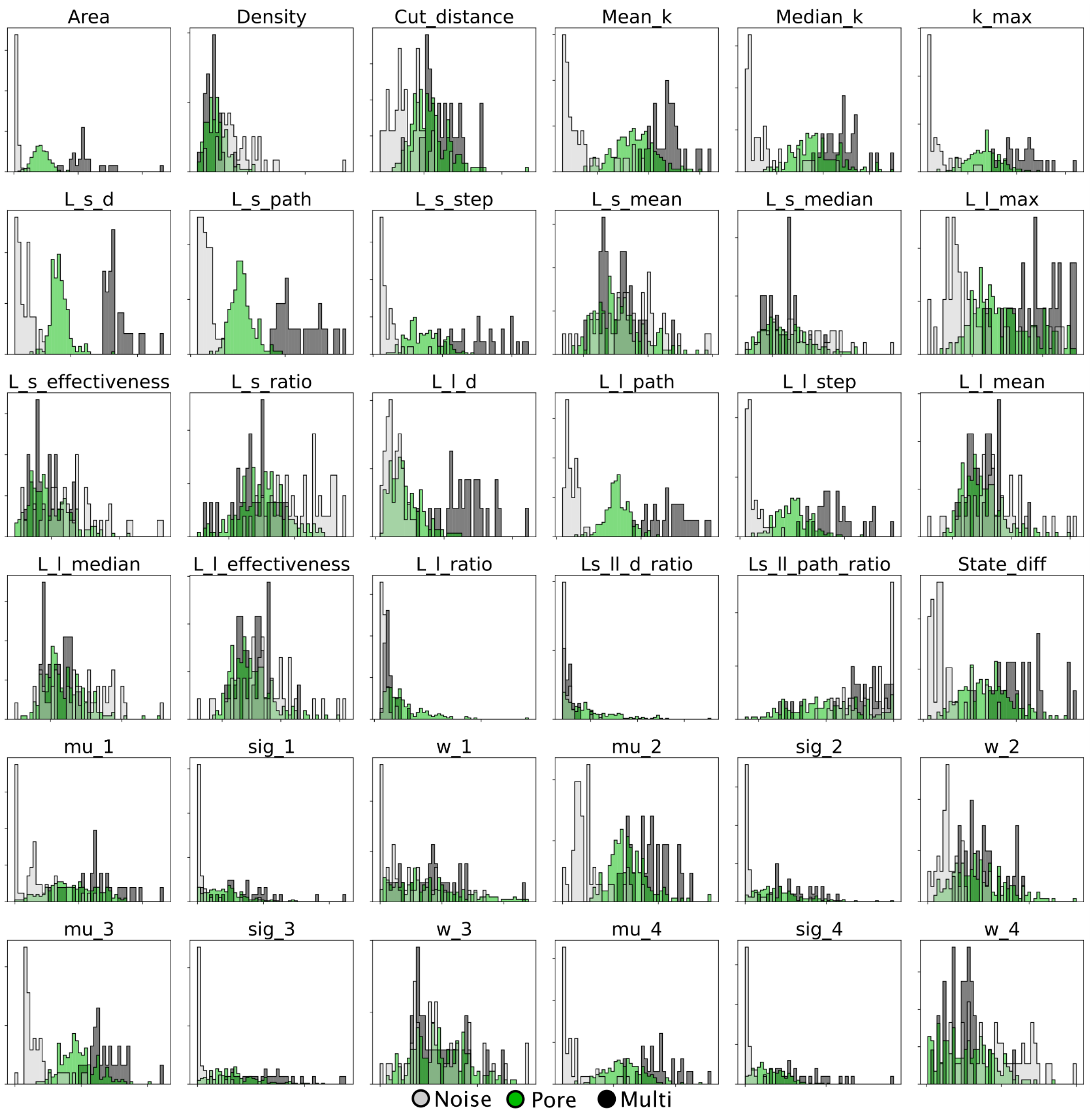


**Supplementary Fig. 11:** All the used spatial features distributions from nuclear pore complex treatment from the NPC-A647 dataset [57]

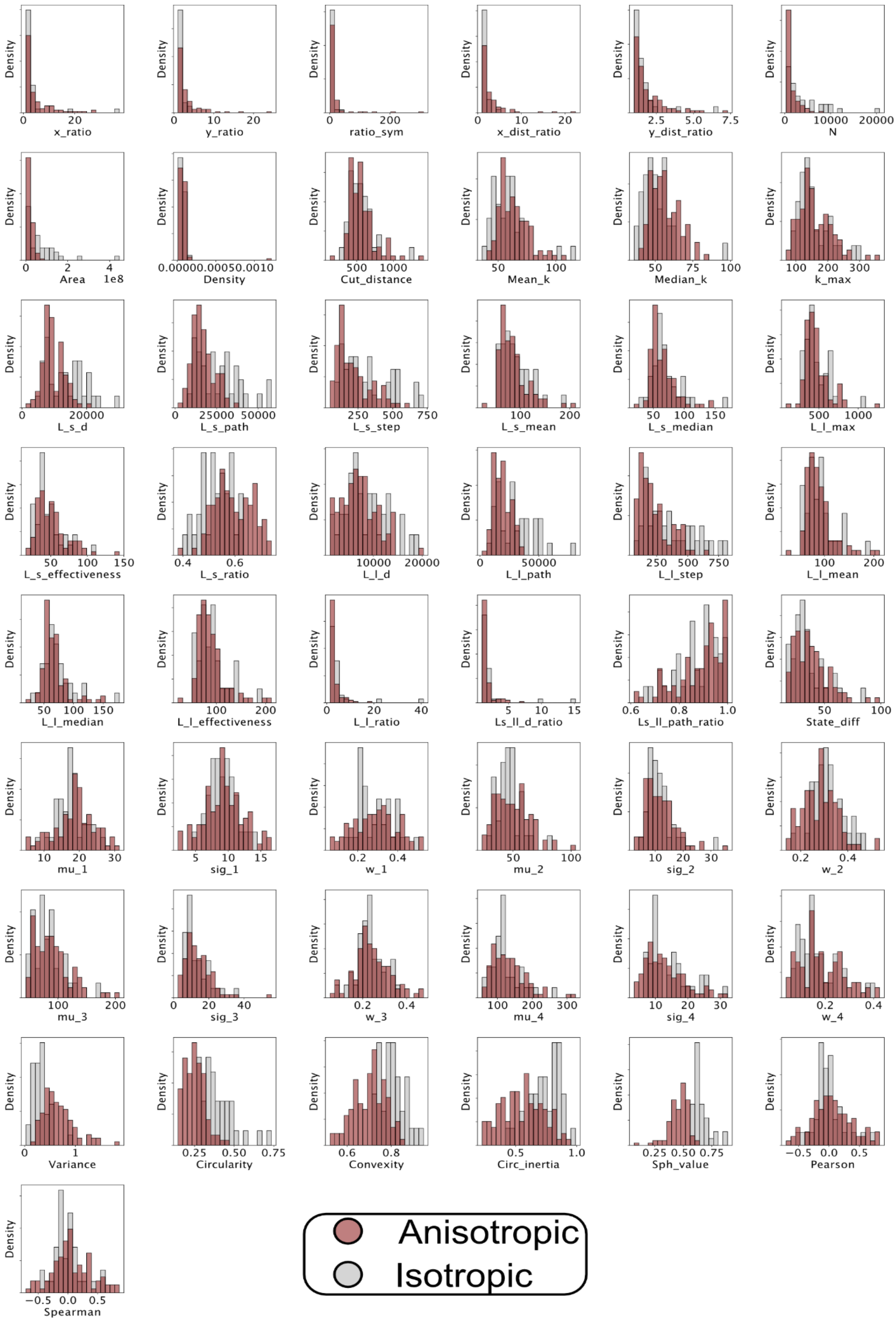


**Supplementary Fig. 12:** Morphology fingerprint feature distributions for anisotropic and isotropic growth type aggregates from the insulin aggregation studies by REPLOM [19].

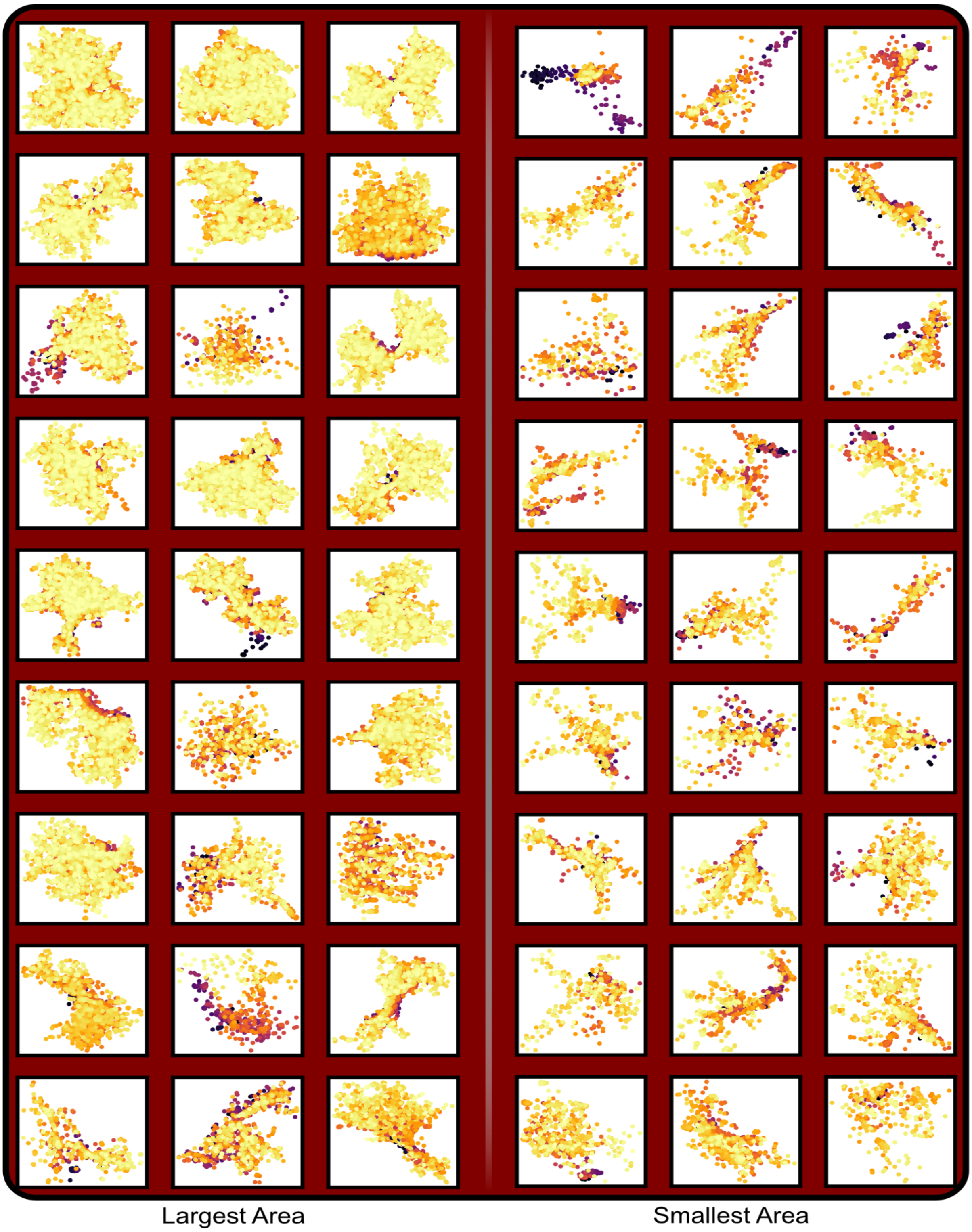


**Supplementary Fig. 13:** The 27 aggregates with the biggest and smallest areas of the anisotropic classified structures from the insulin aggregation studies by REPLOM [19].

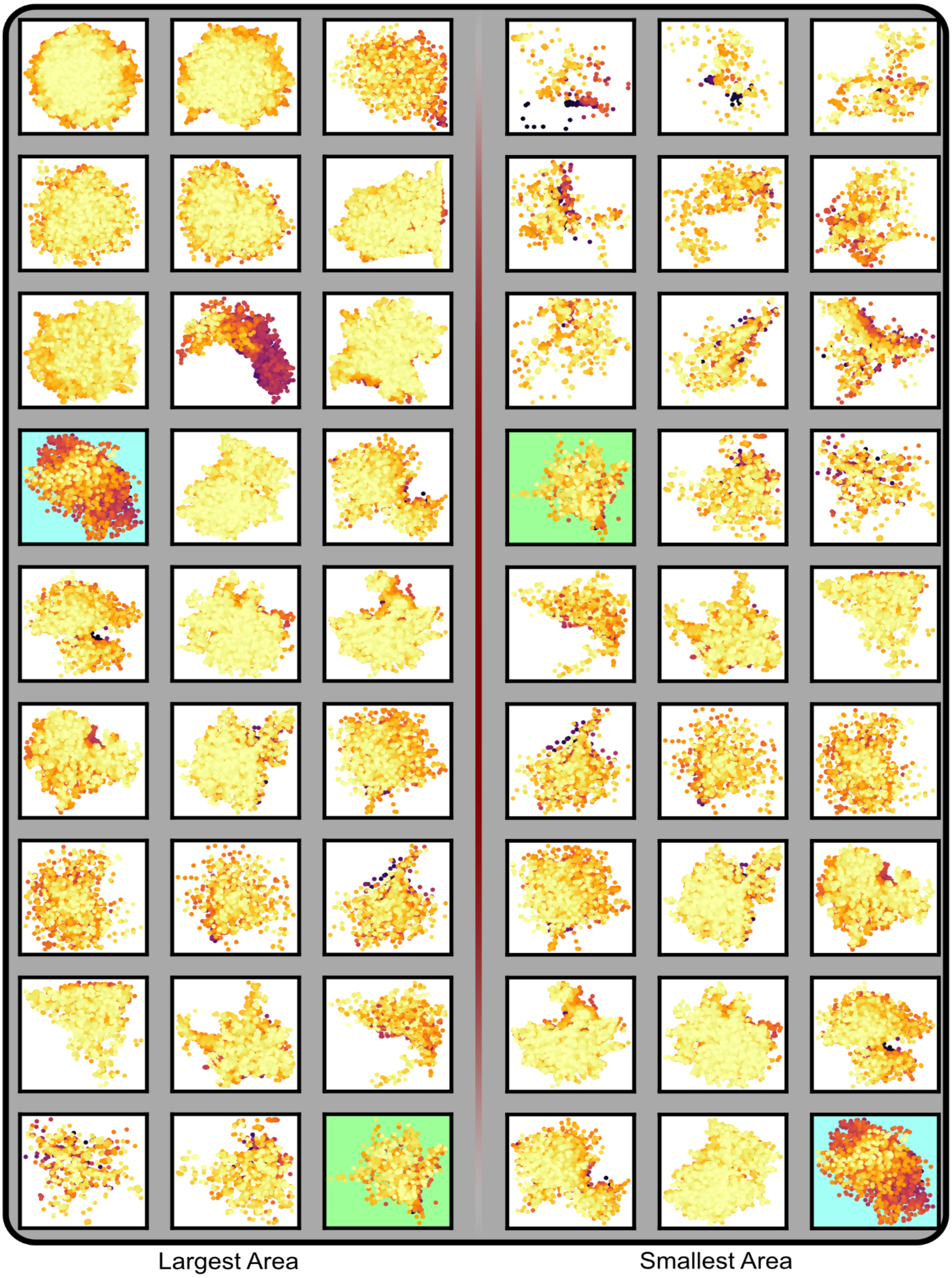


**Supplementary Fig. 14:** The 27 aggregates with biggest and smallest areas of the isotropic classified structures from the insulin aggregation studies by REPLOM [19]. However, as there are only 36 classified Isotropic structures, the green and blue background shows the overlap between the two columns.
